# Rapid degradation of *C. elegans* proteins at single-cell resolution with a synthetic auxin

**DOI:** 10.1101/716837

**Authors:** Michael A. Q. Martinez, Brian A. Kinney, Taylor N. Medwig-Kinney, Guinevere Ashley, James M. Ragle, Londen Johnson, Joseph Aguilera, Christopher M. Hammell, Jordan D. Ward, David Q. Matus

**Affiliations:** Department of Biochemistry and Cell Biology, Stony Brook University, Stony Brook, NY 11794, USA.; Cold Spring Harbor Laboratory, Cold Spring Harbor, NY 11724, USA.; Department of Molecular, Cell, and Developmental Biology, University of California-Santa Cruz, Santa Cruz, CA 95064, USA.

**Keywords:** *C. elegans*, AID system, NAA, Microfluidics, SCF complex, NHR-25

## Abstract

As developmental biologists in the age of genome editing, we now have access to an ever-increasing array of tools to manipulate endogenous gene expression. The auxin-inducible degradation system, allows for spatial and temporal control of protein degradation, functioning through the activity of a hormone-inducible *Arabidopsis* F-box protein, transport inhibitor response 1 (TIR1). In the presence of auxin, TIR1 serves as a substrate recognition component of the E3 ubiquitin ligase complex SKP1-CUL1-F-box (SCF), ubiquitinating auxin-inducible degron (AID)-tagged proteins for proteasomal degradation. Here, we optimize the *Caenorhabditis elegans* AID method, utilizing 1-naphthaleneacetic acid (NAA), an indole-free synthetic analog of the natural auxin indole-3-acetic acid (IAA). We take advantage of the photostability of NAA to demonstrate via quantitative high-resolution microscopy that rapid degradation of target proteins can be detected in single cells within 30 minutes of exposure. Additionally, we show that NAA works robustly in both standard growth media and physiological buffer. We also demonstrate that K-NAA, the water-soluble, potassium salt of NAA, can be combined with microfluidics for targeted protein degradation in *C. elegans* larvae. We provide insight into how the AID system functions in *C. elegans* by determining that TIR1 interacts with *C. elegans* SKR-1/2, CUL-1, and RBX-1 to degrade target proteins. Finally, we present highly penetrant defects from NAA-mediated degradation of the Ftz-F1 nuclear hormone receptor, NHR-25, during *C. elegans* uterine-vulval development. Together, this work provides a conceptual improvement to the AID system for dissecting gene function at the single-cell level during *C. elegans* development.

## INTRODUCTION

*In situ* techniques for targeted protein degradation enable a detailed analysis of developmental events, mechanisms, and functions. RNAi and Cre or Flp-mediated recombination (Qadota *et al*. 2007; Hubbard 2014; Shen *et al*. 2014) allow tissue-specific study of gene products, but the persistence of the target protein following recombination or RNA depletion can delay the manifestation of an otherwise acute phenotype. Several methods have been described recently to enable tissue-specific protein degradation in *Caenorhabditis elegans*, including ZF1 tagging (Armenti *et al*. 2014), a GFP nanobody approach (Wang *et al*. 2017), sortase A (Wu *et al*. 2017), and auxin-mediated degradation (Zhang *et al*. 2015).

The auxin-inducible degradation system allows for rapid and conditional degradation of auxin-inducible degron (AID)-tagged proteins in *C. elegans* as well as in other commonly used model systems including yeast (Nishimura *et al*. 2009), *Drosophila* (Trost *et al*. 2016), zebrafish (Daniel *et al*. 2018), cultured mammalian cells (Nishimura *et al*. 2009; Holland *et al*. 2012; Natsume *et al*. 2016), and mouse oocytes (Camlin and Evans 2019). This protein degradation system relies on the expression of an *Arabidopsis* F-box protein called transport inhibitor response 1 (TIR1). As a substrate-recognition component of the SKP1-CUL1-F-box (SCF) E3 ubiquitin ligase complex, TIR1 carries out its function only in the presence of the hormone auxin. Once bound to auxin, TIR1 targets AID-tagged proteins for ubiquitin-dependent proteasomal degradation (**Figure 1A**).

**Figure 1.**
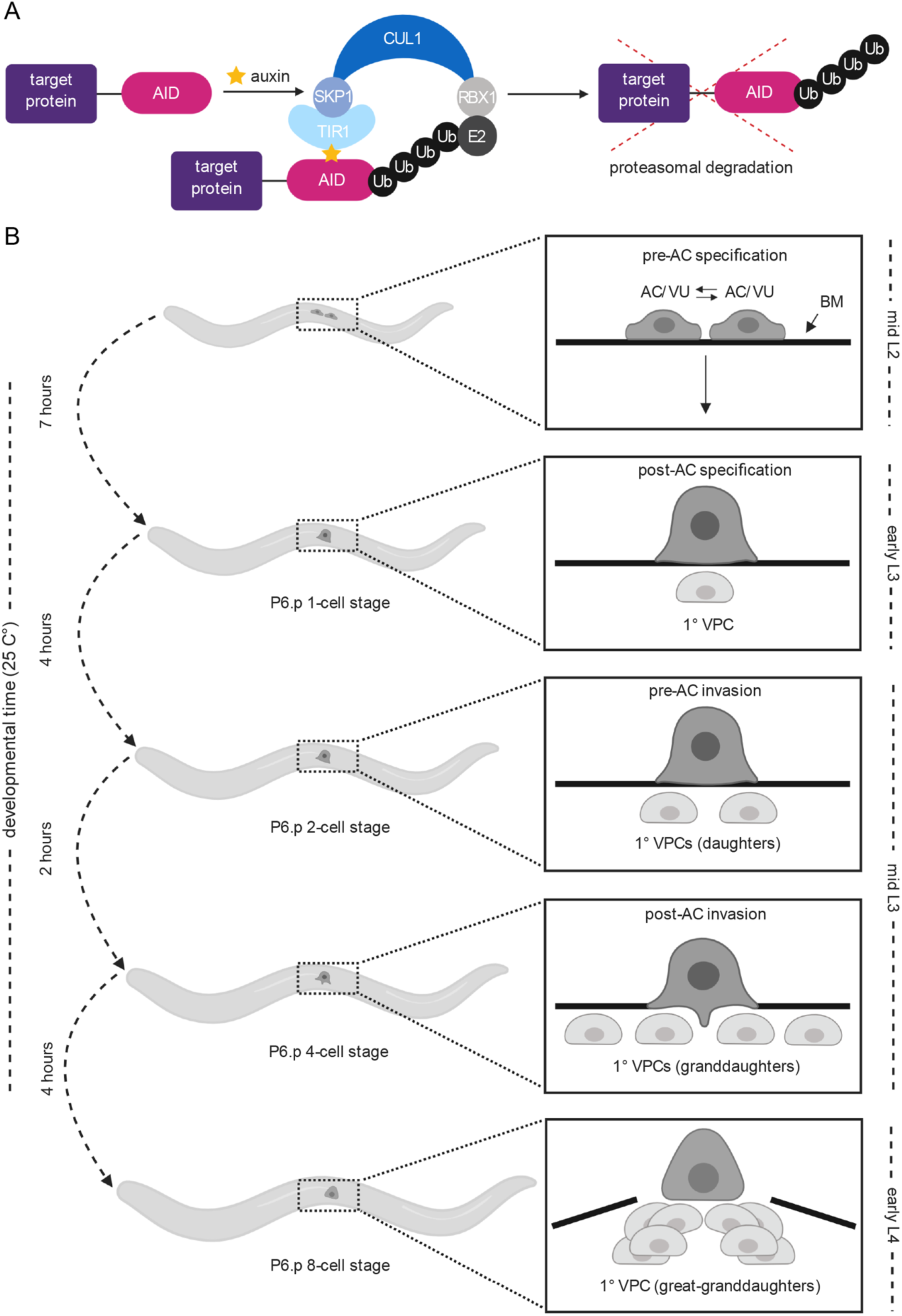
Overview of the auxin-inducible degradation system and *C. elegans* uterine-vulval development. (A) In this system, a target protein is fused to an auxin-inducible degron (AID). Heterologous expression of *Arabidopsis* TIR1 mediates robust auxin-dependent proteasomal degradation of AID-tagged proteins through the SKP1-CUL1-F-box (SCF) E3 ubiquitin ligase complex. (B) Schematic of uterine-vulval morphogenesis during *C. elegans* larval development. In *C. elegans*, AC specification and morphogenesis of uterine-vulval attachment occurs from the mid-L2 through the early L4 stage (Schindler and Sherwood 2013). The AC is specified in a stochastic reciprocal Notch-Delta signaling event in the mid-L2 stage (top panel). Following AC specification, the AC specifies the 1° fate of the underlying vulval precursor cell, P6.p in the early L3 (second panel), which then divides three times to ultimately give rise to eight of the 22 cells of the adult vulva (bottom three panels).

The *C. elegans* version of the AID system is robust and specific with minimal off-target effects (Zhang *et al*. 2015). However, re-evaluation of the system is needed to assess its utility among *C. elegans* researchers conducting microscopy-based single-cell biology within a narrow developmental time frame. Here, we use 1-naphthaleneacetic acid (NAA) and its water-soluble potassium salt analog (K-NAA), indole-free synthetic analogs of the natural auxin indole-3-acetic acid (IAA), to degrade target proteins at single-cell resolution in *C. elegans* larvae in standard growth media and physiological buffer. Given the ability to solubilize K-NAA solely in water or physiological buffer (M9), we also demonstrated rapid degradation kinetics of an AID-tagged transgene in a *C. elegans*-based microfluidics for the first time (Keil *et al*. 2017). Next, we sought to gain insight into which SCF complex members interact with TIR1, identifying *skr-1/2*, *cul-1* and *rbx-1* as putative TIR1 interactors through RNAi depletion experiments. Finally, we demonstrate potent temporal effects on uterine and vulval development following targeted degradation of endogenous NHR-25, the single *C. elegans* homolog of *Drosophila* Ftz-F1 and human SF-1 and LRH-1 (Chen *et al*. 2004; Ward *et al*. 2013). It is our hope that this synthetic auxin analog will be applied at all stages of *C. elegans* development, allowing for precise, rapid degradation of target proteins in a high-resolution and quantitative fashion.

## MATERIALS AND METHODS

### C. elegans strains and culture conditions

Animals were maintained using standard culture conditions at 25°C (Brenner 1974) and were synchronized through alkaline hypochlorite treatment of gravid adults to isolate eggs (Porta-de-la-Riva *et al*. 2012). In the main text and figure legends, we designate linkage to a promoter using a greater than symbol (>) and fusion to a protein using a double colon (::). The following alleles and transgenes were used in this manuscript for experimental purposes: LG I: *kry61[nhr-23::AID]; LG II: ieSi57[eft-3>TIR1::mRuby]; LG IV: ieSi58[eft-3>AID::GFP], syIs49 [zmp-1>GFP]; LG X: wrd10[nhr-25::GFP::AID]*.

### Constructs and microinjection

SapTrap was used to construct the 30xlinker::GFP^SEC^TEV::AID degron::3xFLAG repair template (pJW1747) for generating the knock-in into the 3’ end of the *nhr-25* gene (Schwartz and Jorgensen 2016). DH10β competent *E. coli* cells, made in-house, were used for generating the plasmid. The following reagents were used to assemble the final repair template: pDD379 (backbone with F+E sgRNA), annealed oligos 3482+3483 (sgRNA), pJW1779 (5’ homology arm), 3’ homology arm PCR product, pJW1347 (30x linker for CT slot), pDD372 (GFP for FP slot), pDD363 (SEC with LoxP sites), and pJW1759 (TEV::AID degron::3xFLAG for NT slot).

The pJW1747 repair template was purified using the Invitrogen PureLink HQ Mini Plasmid DNA Purification Kit (K210001). The optional wash step in the protocol using a 4 M guanidine-HCl + 40% isopropanol solution is highly recommended, as excluding it dramatically reduced injection efficiency in our hands. N2 animals were injected with a mix consisting of 10 ng/µl of pJW1747, 50 ng/µl of pDD121 (Cas9 vector), and co-injection markers (10 ng/µl pGH8, 5 ng/µl pCFJ104, 2.5 ng/µl pCFJ90) as previously described (Frøkjær-Jensen *et al*. 2012; Dickinson *et al*. 2013, 2015). Knock-ins were isolated as previously described (Dickinson *et al*. 2015). Each knock-in junction was verified via PCR using a primer that bound outside the homology arm paired with a primer binding within pJW1747. The knock-in was backcrossed five times against wild-type N2 animals to produce JDW58. The SEC was then excised by heat-shock (Dickinson *et al*. 2015) to produce JDW59; the knock-in sequence was re-confirmed by PCR amplification and sequencing, using the oligos flanking the homology arms. JDW58 was crossed to CA1200 (*eft-3>TIR1::mRuby*) to generate JDW70. The SEC was then excised (Dickinson *et al*. 2015) to produce JDW71.

pDD121, pDD363, pDD372, and pDD379 (Dickinson *et al*. 2018) were gifts from Bob Goldstein (Addgene plasmid numbers are 91833, 91829, 91824, and 91834, respectively). pJW1347 and pJW1759 will be deposited into Addgene’s repository and are also available upon request. pJW1347 and pJW1759 were generated by TOPO blunt cloning of PCR products. pJW1779 was generated by Gateway cloning into pDONR221 (Invitrogen). Oligo sequences used to generate these plasmids, the sgRNA, the 3’ homology arm, and for genotyping are in the Reagent Table.

### Auxin experiments

For all auxin experiments, synchronized L1 larval stage animals were first transferred to standard nematode growth media (NGM) agar plates seeded with *E. coli* OP50 and then transferred at the P6.p 2-cell stage (mid-L3 stage) to either OP50-seeded NGM agar plates treated with IAA, NAA, or K-NAA, or M9 buffer treated with NAA in the absence of bacteria, NAA plus *E. coli* NA22, or K-NAA plus *E. coli* NA22.

For IAA and NAA experiments on plates, a 250 mM stock solution in 95% ethanol was prepared using powder IAA purchased from Alfa Aesar (A10556) and powder NAA purchased from Sigma-Aldrich (317918) and stored at −20°C. IAA and NAA were then diluted into the NGM agar (cooled to approximately 50°C) at the time of pouring plates. Fresh OP50 was used to seed plates. For control experiments, 0.25% ethanol was used as described previously (Zhang *et al*. 2015). For K-NAA experiments on plates, a 250 mM stock solution in deionized water was prepared using powder K-NAA purchased from PhytoTechnology Laboratories (N610) and stored at 4°C. For control experiments, OP50-seeded NGM agar plates were used. Prior to each NAA experiment in M9 buffer, a fresh 1 mM (pH of 7.22) or 4 mM solution (pH of 8.14) in M9 buffer was prepared using 5.4 mM NAA purchased in liquid form from Sigma-Aldrich (N1641). These pH levels are well within the tolerance range of *C. elegans* for pH (Khanna *et al*. 1997). M9 buffer alone (pH of 7.13) was used as a control. A detailed protocol for liquid-based NAA-mediated degradation can be found in File S1. For experiments conducted in the microfluidic platform, a 4 mM NAA or K-NAA solution in M9 buffer containing *E. coli* NA22 was prepared and stored at 4°C for up to 2 weeks. M9 buffer containing NA22 was used as a control. See File S2 for a detailed protocol describing the preparation of media for the microfluidic device.

### Brood size and viability assays

Brood size and viability assays were performed as described (Zhang *et al*. 2015). Briefly, L4 hermaphrodites were picked onto individual MYOB plates containing 0% ethanol (K-NAA control), 0.25% ethanol (IAA control), 4 mM K-NAA, or 4 mM IAA. Animals were then transferred to new plates daily over 4 days. The eggs laid on each plate were counted after removing the parent and viable progeny were quantified when the F1 reached L4 or adult stages (2-3 days post egg-laying). At this point, we also scored for dead eggs. Brood size is the sum of live progeny and dead eggs. Percent embryonic lethality was determined by dividing dead eggs by total eggs laid.

### RNAi experiments

RNAi targeting *cul-1* was constructed by cloning 997 bp of synthetic DNA based on its cDNA sequence available on WormBase (wormbase.org) into the highly efficient T444T RNAi vector (Sturm *et al*. 2018). The synthetic DNA was generated by Integrated DNA Technologies (IDT) as a gBlock gene fragment and cloned into the BglII/SalI restriction digested T444T vector using the NEBuilder HiFi DNA Assembly Master Mix (E2621). RNAi feeding strains silencing *skr-1/2*, *skr-7*, *skr-10*, and *rbx-1* were obtained from the Vidal RNAi library (Rual *et al*. 2004).

### Scoring defects in anchor cell (AC) specification

Synchronized L1 stage *nhr-25::GFP::AID; eft-3>*TIR1::mRuby animals were plated onto NGM agar plates containing either control or 4 mM NAA and grown for 24 hours at 25°C until the early L3 stage (P6.p 1-cell stage), after the normal time of AC specification. Images were acquired as specified below to score for the presence or absence of an AC, visualized by characteristic morphology using DIC optics.

### Scoring vulva precursor cell (VPC) arrest

Synchronized L1 stage *nhr-25::GFP::AID; eft-3>*TIR1::mRuby animals were plated onto OP50 NGM agar plates and allowed to grow until the P6.p 1-cell stage. Animals were then washed off plates with M9 and transferred onto NGM agar plates containing either control or 4 mM NAA and grown at 25°C until the mid-L3 stage, after the normal time of P6.p cell division. Images were acquired as specified below to score for P6.p divisions using DIC optics. Remaining animals were scored for plate level adult phenotypes approximately 24 hours later.

### Image acquisition

Images were acquired using a Hamamatsu Orca EMCCD camera and a Borealis-modified Yokagawa CSU-10 spinning disk confocal microscope (Nobska Imaging, Inc.) with a Plan-APOCHROMAT x 100/1.4 oil DIC objective controlled by MetaMorph software (version: 7.8.12.0). Animals were anesthetized on 5% agarose pads containing 10 mM sodium azide and secured with a coverslip. Imaging on the microfluidic device was performed on a Zeiss AXIO Observer.Z7 inverted microscope using a 40X glycerol immersion objective and DIC and GFP filters controlled by ZEN software (version 2.5). Images were captured using a Hamamatsu C11440 digital camera. For scoring plate level phenotypes, images were acquired using a Moticam CMOS (Motic) camera attached to a Zeiss dissecting microscope.

### Image processing and analyses

All acquired images were processed using Fiji software (version: 2.0.0-rc-69/1.52p) (Schindelin *et al*. 2012). To quantify AC- or VPC-specific degradation of AID::GFP, images were captured at the P6.p 2-cell stage and 4-cell stage (mid-L3 stage) at time points 0, 30, 60, 90, and 120 minutes in the absence or presence of auxin. Expression of *eft-3>*AID::GFP was quantified by measuring the mean fluorescence intensity (MFI) of ACs and VPCs subtracted by the MFI of a background region in the image to account for camera noise. Cells were outlined using the freehand selection tool in Fiji. Data were normalized by dividing the MFI in treated or untreated animals at time points 30, 60, 90, and 120 minutes by the average MFI in untreated animals at 0 minutes. For experiments utilizing RNAi, only ACs were measured due to the variable sensitivity of VPCs to RNAi (Bourdages *et al*. 2014; Matus *et al*. 2014). To quantify AC-specific degradation of AID::GFP in animals fed RNAi overnight, images were captured at the P6.p 2-cell stage before auxin treatment and 60 minutes post-treatment. *eft-3>*AID::GFP expression in the AC was quantified as described above. Data were normalized by dividing the MFI in auxin treated animals by the average MFI in untreated animals. To analyze *nhr-25::GFP::AID* degradation, GFP levels were quantified by measuring the MFI in individual GFP-expressing nuclei in the AC/VU, AC, or VPCs subtracted by the MFI of a background region in the image to account for background noise. Nuclei were outlined using the threshold tool in Fiji or for animals with no detectable GFP signal, the corresponding DIC image was utilized to identify the nucleus. Images of L3 larvae were captured in a *C. elegans* larvae-specific microfluidic device (Keil *et al*. 2017). To quantify AID::GFP degradation, animals were loaded into the microfluidic chamber and fed NA22 bacteria. Images were captured at time points 0, 30, 60, 90, and 120 minutes with or without auxin. Here, *eft-3>*AID::GFP expression was quantified by measuring the MFI in whole animals subtracted by the MFI of a background region in the image to account for background noise. Whole animals were outlined using the freehand selection tool in Fiji. Data were normalized by dividing the MFI in treated or untreated animals at time points 30, 60, 90, and 120 minutes by the average MFI in untreated animals at timepoints 30, 60, 90, and 120 minutes respectively to account for photobleaching from imaging the same animal. Cartoons were created with BioRender (biorender.com) and ChemDraw software (version: 18.0). Graphs were generated using Prism software (version: 8.1.2). Figures were compiled using Adobe Photoshop (version: 20.0.6) and Illustrator (version: 23.0.26).

### Statistical analyses

A power analysis was performed to determine the sample size (*n*) needed per experiment to achieve a power level of 0.80 or greater (Cohen 1992; Pollard *et al*. 2019). Statistical significance was determined using either a two-tailed unpaired Student’s t-test or Mann Whitney U test. *P* < 0.05 was considered statistically significant. The figure legends specify when error bars represent the standard deviation (SD) or interquartile range (IQR).

### Data availability

Supplemental data and key reagents can be found at Figshare. Worm strains CA1202, CA1204, and PS3239 are available to order from the *Caenorhabditis* Genetics Center. All other strains are available upon request. The data that support the findings of this study are available upon reasonable request.

## RESULTS AND DISCUSSION

### NAA is a synthetic alternative to the natural auxin IAA

Given the recent advances in CRISPR/Cas9-genome editing technology (Dickinson and Goldstein 2016; Dokshin *et al*. 2018), the auxin-inducible degron (AID) with or without a fluorescent reporter (e.g., GFP or its derivatives) can be inserted into a genomic locus of interest (Röth *et al*. 2019). Though this technology can be applied with ease, there are certain limitations that exist with the use of the natural auxin indole-3-acetic acid (IAA), including its limited solubility in water. While levels of the ethanol solvent used to dissolve IAA (0.25–1.52%) are well below the threshold for causing a physiologic response (Morgan and Sedensky 1995; Kwon *et al*. 2004), higher percentages of ethanol (7%) have been shown to cause rapid changes in *C. elegans* gene expression (Kwon *et al*. 2004). A potentially more problematic limitation of IAA for live-cell imaging-based applications is cytotoxicity related to excitation with UV and blue light. Specifically, IAA has been shown in yeast (Papagiannakis *et al*. 2017) and mouse oocytes (Camlin and Evans 2019) to cause cytotoxicity, likely due to acceleration of the oxidative decarboxylation of IAA to methylene-oxindole (Srivastava 2002). In yeast, IAA exposure during live-cell imaging suppressed cell proliferation (Papagiannakis *et al*. 2017) and mammalian oocytes failed to complete meiotic maturation (Camlin and Evans 2019). In both systems, the use of a synthetic auxin, 1-naphthaleneacetic acid (NAA), rescued these cytotoxic responses. For these reasons, we chose to examine whether NAA would also function in *C. elegans* to degrade AID-tagged proteins in the presence of TIR1. Ultimately, we wished to evaluate AID-mediated degradation (**Figure 1A**) in single cells and tissues using live-cell imaging. Thus, we determined the kinetics of protein degradation using spinning disk confocal microscopy, rather than using low-magnification microscopy and Western blot analysis to measure protein loss in whole animals, as performed previously (Zhang *et al*. 2015). We chose to focus primarily on the L3 stage of post-embryonic development due to many of the dynamic cellular behaviors occurring over relatively short time scales (within minutes to hours) in this developmental window, including uterine-vulval attachment and vulval morphogenesis (**Figure 1B**) (Gupta *et al*. 2012).

To analyze the kinetics of AID-mediated degradation in the uterine anchor cell (AC) and underlying vulva precursor cells (VPCs) (**Figure 1B**), we utilized a previously published strain expressing AID::GFP and TIR1::mRuby under the same ubiquitously expressed *eft-3* promoter (Zhang *et al*. 2015). The single-cell abundance of AID::GFP was measured over time in mid-L3 stage animals exposed to different concentrations of auxin incorporated into standard *C. elegans* solid culture media (**Figure 2A**). In addition to testing the natural auxin IAA, also tested whether it was possible to perform auxin-inducible degradation in the AC and VPCs using the synthetic auxin analog NAA (**Figure 2B**). In the presence of ≥ 1 mM IAA or NAA, AID::GFP abundance in the AC and VPCs was reduced by approximately 80% of its initial level within 30 minutes (**Figure 2C-D**). Within 60 minutes, AID::GFP was virtually undetectable (**Figure 2C-D**). These results indicate that NAA can serve as a viable substitute to IAA. These results indicate that NAA can serve as a viable substitute for IAA. Similar to (Zhang *et al*. 2015), we observed that growth on IAA resulted in similar brood sizes compared to control (**Table S1**). Growth on the potassium salt of NAA (K-NAA) resulted in similar brood sizes compared to control, and also produced a modest but significant (*P* = 0.02) reduction in embryonic lethality compared to IAA treatment (**Table S1**). Consistent with this result, higher levels of toxicity was observed when using IAA over NAA in studies investigating circadian rhythm biology in *Drosophila* (Chen *et al*. 2018). Zhang *et al*. (2015) reported inhibited bacterial growth at high concentrations of auxin. Compared to bacterial growth on IAA and NAA, we observed more robust OP50 growth on K-NAA with no trade-off in degradation rate (**Figure S1A).** Together, these results demonstrate that NAA is a viable alternative to IAA for targeted protein degradation in *C. elegans* larvae.

**Figure 2.**
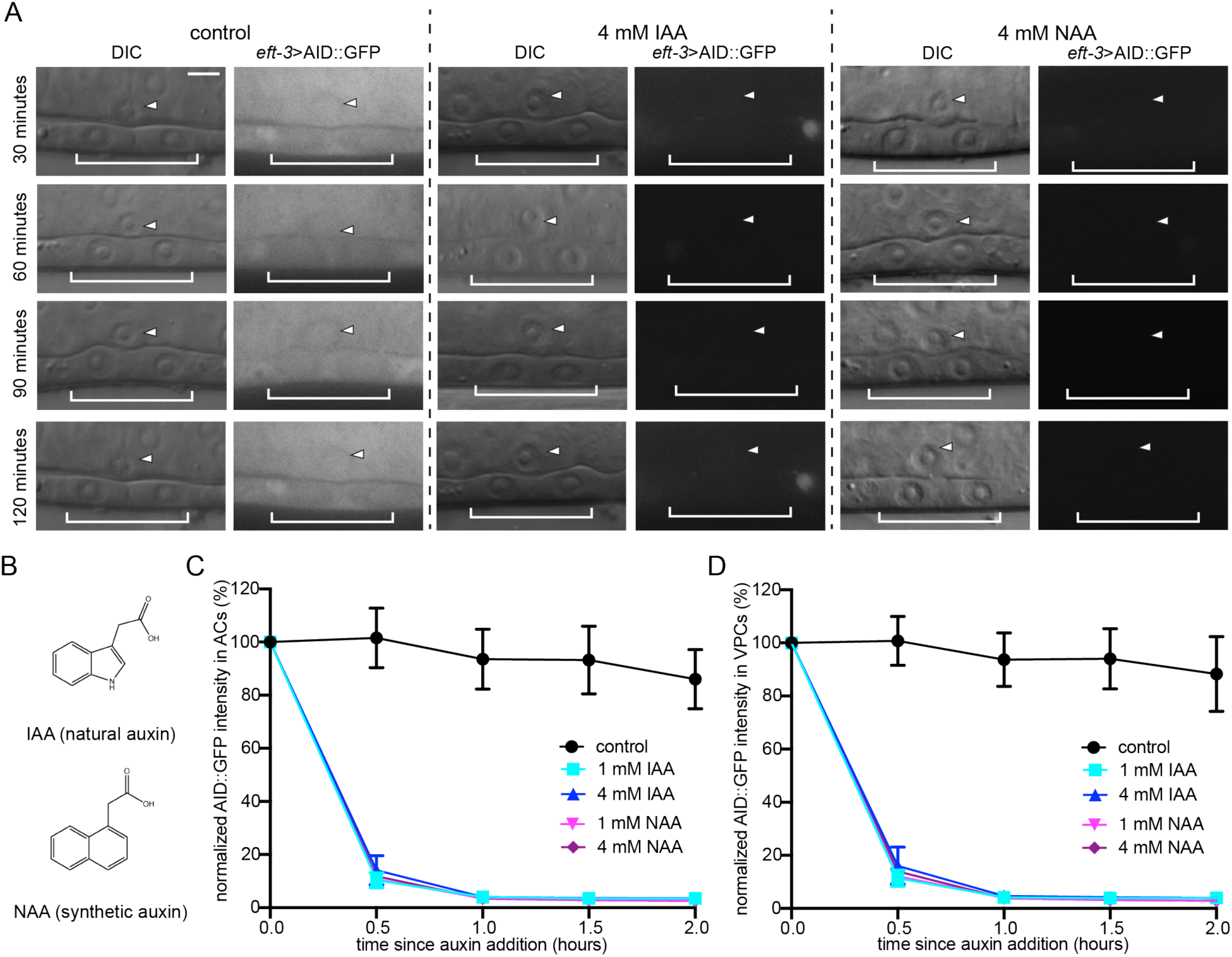
Comparison of IAA- and NAA-mediated degradation in the *C. elegans* AC and VPCs. (A) DIC and corresponding GFP images of ACs (arrowheads) and underlying 1° fated VPCs (brackets) from mid-L3 stage animals at the P6.p 2-cell stage. Animals expressing AID::GFP and TIR1::mRuby under the same *eft-3* promoter were treated with natural auxin indole-3-acetic acid (IAA) and synthetic auxin 1-naphthaleneacetic acid (NAA) in NGM agar containing OP50. (B) Chemical structure of IAA and NAA. (C, D) Rates of degradation determined by quantifying AID::GFP in (C) ACs and (D) VPCs following auxin treatment. Data presented as the mean±SD (*n* ≥ 30 animals examined for each time point).

### NAA is soluble in physiological buffer

To compare degradation kinetics on plate-based growth to depletion in liquid culture during imaging, we measured AID::GFP abundance in the AC and VPCs in mid-L3 stage animals exposed to different concentrations of NAA solubilized in M9 buffer (**Figure 3A-D**). In the presence of ≥ 1 mM liquid NAA, AID::GFP in the AC and VPCs was reduced by 80%, as compared to initial levels, within 30 minutes and was nearly undetectable within 60 minutes (**Figure 3B-D**). These results show that NAA can induce auxin-dependent degradation in liquid culture in *C. elegans*, reducing the need to rear animals on auxin plates and transfer to slides for imaging. This finding raises the possibility of depleting proteins and imaging the resulting developmental consequences at single-cell resolution.

**Figure 3.**
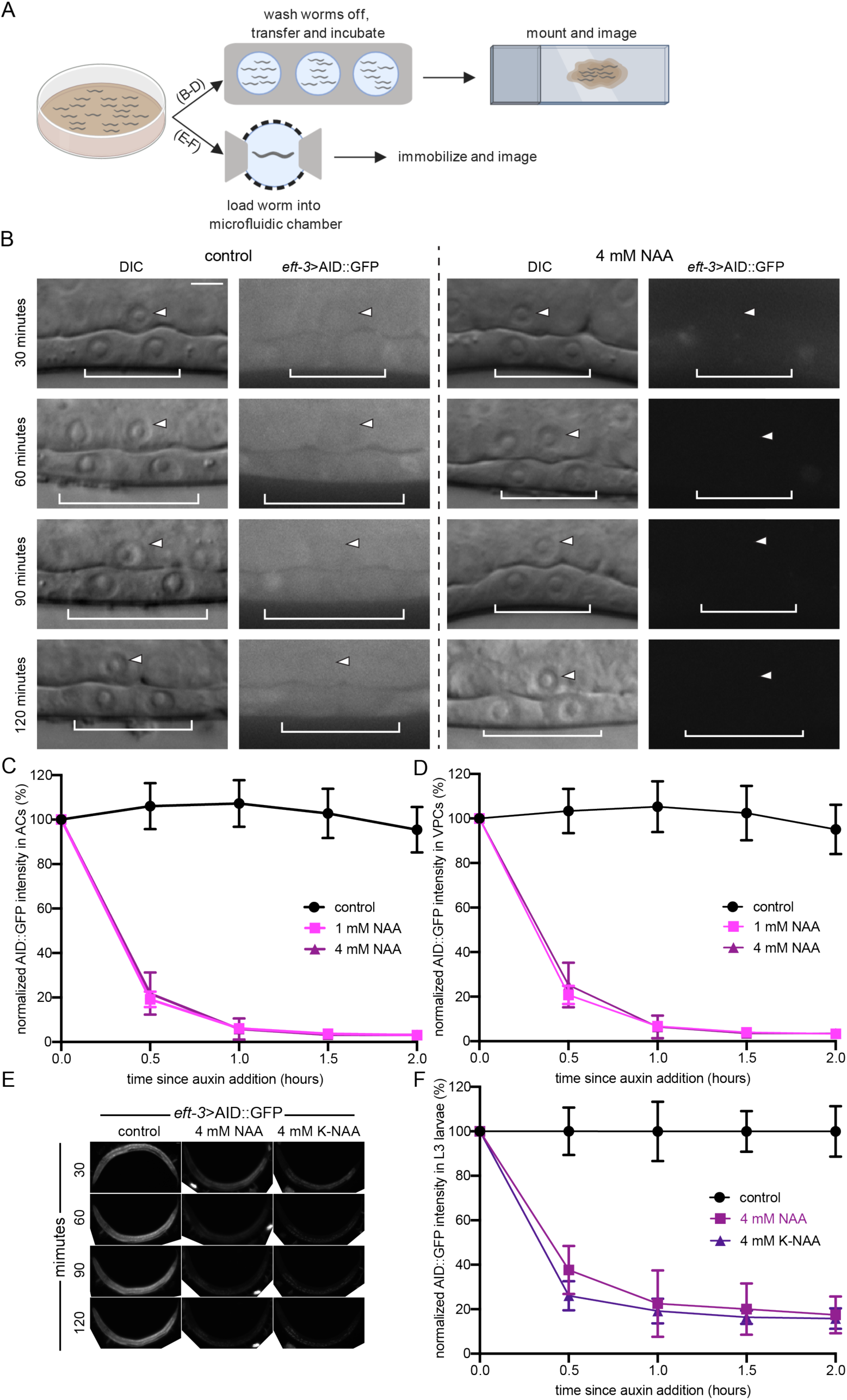
Solubility of NAA/K-NAA in physiological buffer enhances utility. (A) Schematic representation of the liquid NAA-based degradation protocol for use in high-resolution microscopy or microfluidics-based approaches. (B) DIC and corresponding GFP images of ACs (arrowheads) and underlying VPCs (brackets) from mid-L3 stage animals at the P6.p 2-cell stage. Animals expressing AID::GFP and TIR1::mRuby under the same *eft-3* promoter were treated with NAA in M9. (C, D) Rates of degradation were determined by quantifying AID::GFP in (C) ACs and (D) VPCs following auxin treatment. Data presented as the mean±SD (*n* ≥ 30 animals examined for each time point). (E) Images of AID::GFP expression from mid-L3 stage animals in control conditions (M9 buffer containing NA22 only, left) or conditions where a 4 mM NAA (middle) or K-NAA (right) solution in M9 buffer containing NA22 was perfused through the microfluidic chamber for the time indicated (Keil et al. 2017). Anterior is left and ventral is down. (F) Rates of degradation were determined by quantifying AID::GFP in whole animals following auxin treatment. Data presented as the mean±SD (*n* ≥ 4 animals examined for each time point).

### K-NAA is an option for C. elegans researchers employing microfluidics

The ability to easily solubilize NAA in physiological buffer raises the possibility of performing protein degradation experiments paired with microfluidics where individual animals can be imaged over long periods at cellular resolution. *C. elegans* lifespan and behavioral assays can involve, subtle phenotypes sensitive to environmental perturbations. Accordingly, the low level of ethanol present in IAA plates is not optimal for these types of assays. The reduced bacterial growth on ethanol and IAA could also affect nutrition (**Figure S1A**) (Cabreiro *et al*. 2013), making water soluble K-NAA an attractive alternative. To compare degradation kinetics between NAA and K-NAA, we time-lapsed L3 stage animals using a microfluidic device optimized for long-term imaging of *C. elegans* larvae (Keil *et al*. 2017), assessing depletion of ubiquitously expressed AID::GFP in trapped animals (**Figure 3A**). At the L3 stage, animals were loaded into the microfluidic chamber in M9 and flushed with a mixture of M9, 4 mM NAA or K-NAA, and NA22 *E. coli* as a bacterial food source (Keil *et al*. 2017). The animals were imaged every 30 minutes for 2 hours. During image acquisition, animals were temporarily immobilized by manually increasing the negative pressure on the compression layer of the device (Keil *et al*. 2017). Between timepoints, animals were allowed to move and feed freely in 4 mM NAA or K-NAA combined with NA22 in M9. Although degradation kinetics were slower than those observed in NAA solubilized in M9 alone (**Figure 3B-D**) or NGM plates containing NAA or K-NAA (**Figure S1B-C**), we still observed approximately 60–70% reduction of AID::GFP expression within the first 30 minutes of NAA or K-NAA exposure, and nearly 80% depletion of whole animal AID::GFP within 1 hour (**Figure 3E-F**). Our results may be under-representing the overall loss of AID::GFP as we did not account for gut autofluorescence in our quantification of fluorescence intensity in whole animals (Teuscher and Ewald 2018). Nonetheless, our results demonstrate that AID-tagged proteins can be depleted in a microfluidic platform which, when combined with long-term high-resolution imaging, provides a powerful tool for studying post-embryonic *C. elegans* development at cellular resolution.

### The AID system functions through specific components of the C. elegans SCF complex to degrade target proteins

Our work demonstrates that the AID system functions rapidly to degrade target proteins in *C. elegans* as described previously (Zhang *et al*. 2015). As a heterologous system, researchers have shown in yeast that *Arabidopsis* TIR1 interacts with the *S. cerevisiae* Cul1 homolog, Cdc53 (Nishimura *et al*. 2009). Whether *Arabidopsis* TIR1 also functions through *C. elegans* proteins homologous to yeast SCF proteins is unknown. Thus, to examine interactions between TIR1 and SCF complex proteins in *C. elegans*, we used RNAi technology directed against components of the SCF complex and quantified AID::GFP in the presence and absence of NAA (**Figure 4 and Figures S2-S3**). This experiment was designed to provide insight into the mechanism through which the AID system depletes target proteins in *C. elegans* and as an intersectional proof-of-concept test of combining auxin-based depletion with a RNAi feeding approach. Briefly, the SCF complex consists of three components: SKP1, CUL1, and RBX1. In contrast to yeast and humans, which contain only one functional SKP1 protein, the scaffold protein CUL-1 is known to interact with eight of the Skp1-related adaptor proteins in *C. elegans*, including SKR-1, −2, −3, −4, −7, −8, −9 and −10 (Nayak *et al*. 2002; Yamanaka *et al*. 2002). We first perturbed *cul-1* expression. To deplete CUL-1, we generated a new RNAi construct targeting *cul-1* in the upgraded T444T RNAi targeting vector (Sturm *et al*. 2018). Notably, this vector contains T7 terminator sequences, which prevents non-specific RNA fragments from being synthesized from the vector backbone (Sturm *et al*. 2018). This vector modification increases the efficiency of mRNA silencing over the original L4440 vector (Sturm *et al*. 2018).

**Figure 4.**
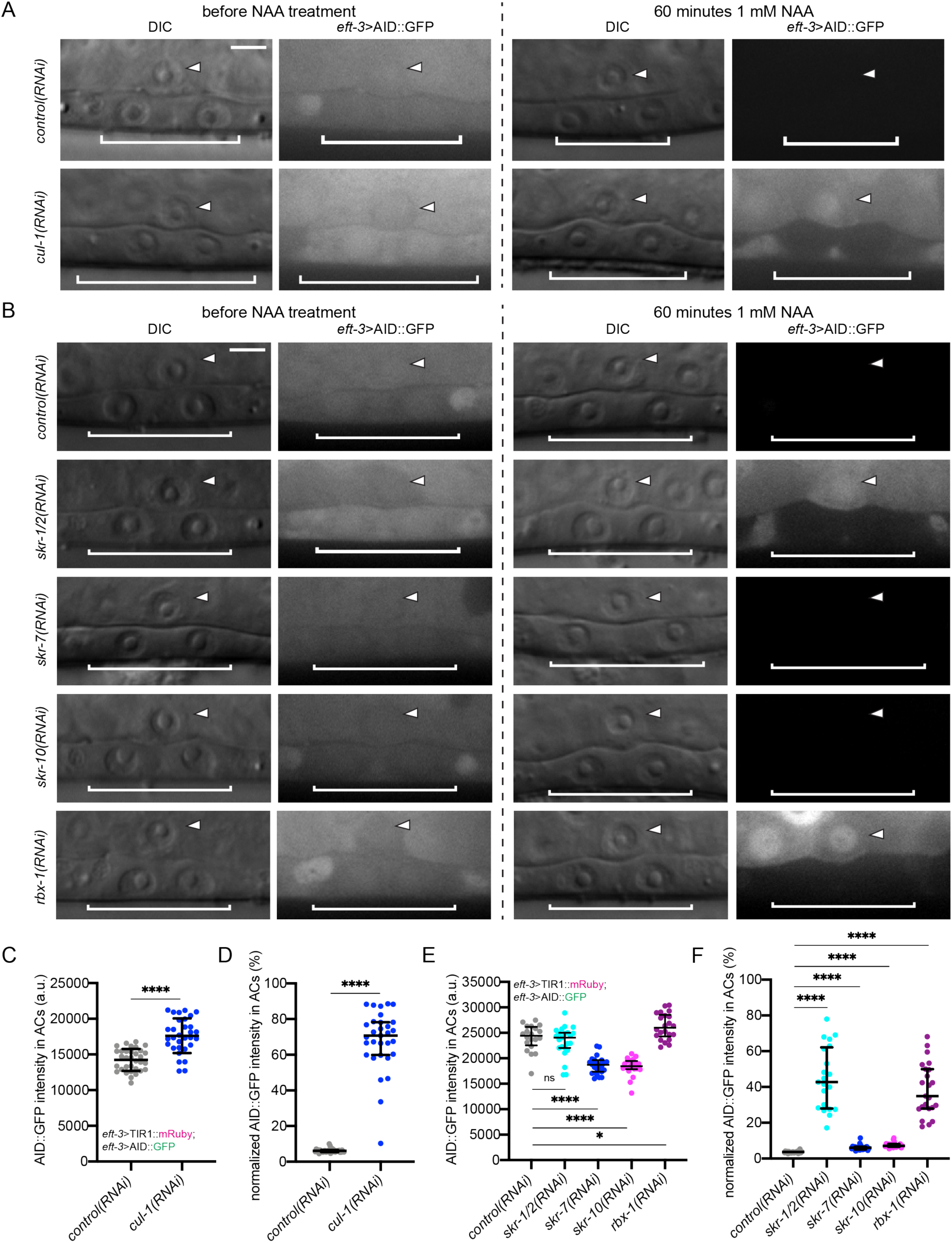
Suppression of SCF complex member expression inhibits TIR1-dependent degradation in the *C. elegans* AC. (A-B) DIC and corresponding GFP images of ACs (arrowheads) and underlying VPCs (brackets) from mid-L3 stage animals at the P6.p 2-cell stage. Animals expressing AID::GFP and TIR1::mRuby under the same *eft-3* promoter were treated with (A) *cul-1(RNAi)* and (B) *skr-1/2, skr-7, skr-10* and *rbx-1(RNAi)*. (C) Quantification of AID::GFP in ACs following *cul-1(RNAi)* treatment. Data presented as the mean±SD (*n* ≥ 30 animals examined for each, and *P* < 0.0001 by a Student’s t-test). (D) Quantification of AID::GFP in ACs following treatment with NAA. Data presented as the median+IQR (*n* ≥ 30 animals examined for each, and *P* < 0.0001 by a Mann Whitney U test). (E) Quantification of AID::GFP in ACs following RNAi knockdown of Skp1-related (*skr*) genes and *rbx-1*. Data presented as the median+IQR (*n* ≥ 20 animals examined for each, and *P* values examined by a Mann Whitney U test). (F) Quantification of AID::GFP in ACs following treatment with NAA. Data presented as the median+IQR (*n* ≥ 21 animals examined for each, and *P* values examined by a Mann Whitney U test).

We hypothesized that depleting CUL-1 would strongly interfere with the proteasomal machinery and thus protein turnover. To assess the abundance of AID::GFP in the *C. elegans* AC, we treated animals with either *control(RNAi)* or *cul-1(RNAi)*. As before, we made use of a strain expressing AID::GFP and TIR1::mRuby from an *eft-3* driver (Zhang *et al*. 2015), and we examined animals at the P6.p 2-cell stage (**Figure 4A**). RNAi knockdown of *cul-1* resulted in a modest but statistically significant increase in the abundance of AID::GFP in the AC compared to *control(RNAi)* treatment (+19%, *n* = 31 and 33, respectively, *P* < 0.0001) (**Figure 4C**). This result suggests that there is TIR-dependent, auxin-independent depletion of AID::GFP, similar to reports in other systems (Morawska and Ulrich 2013; Nishimura and Fukagawa 2017; Zasadzińska *et al*. 2018). To further test this notion, we assessed GFP abundance in the AC in animals lacking TIR1::mRuby. AID::GFP and GFP were driven by *eft-3* and *zmp-1* promoters, respectively, and we performed the experiment on animals at the P6.p 2-cell stage. We did not assess AID::GFP abundance in the VPCs due to the variable sensitivity of this tissue to RNAi (compare **Figure 4A and S2**) (Bourdages *et al*. 2014; Matus *et al*. 2014). Depletion of CUL-1 in animals expressing AID::GFP (−1.8%, *n* = 29 for both treatments, *P* = 0.1168; **Figure S3A-B**) or GFP (−2.4%, *n* = 20 for both RNAi treatments, *P* = 0.7682; **Figure S3C-D**) resulted in a slight decrease in GFP abundance, though it was not statistically significant in either case compared to treatment with *control(RNAi)*. The modest decrease in protein abundance in animals lacking TIR1::mRuby suggests that knockdown of *cul-1* might mildly perturb protein homeostasis, but TIR1-mediated proteosomal degradation of AID-tagged proteins independent of auxin exposure requires endogenous levels of CUL-1 to function robustly.

Next, we tested whether depletion of *cul-1* would inactivate AID-mediated protein degradation. We fed synchronized L1 stage animals with RNAi targeting *cul-1*, and then treated animals at the P6.p 2-cell stage with 1 mM NAA for 60 minutes before quantifying AID::GFP degradation in the AC (**Figure 4A**). For *control(RNAi)*-treated animals, the abundance of AID::GFP in the AC was nearly undetectable within 60 minutes (−94%, *n* = 33; **Figure 4D**). Remarkably, the abundance of AID::GFP in the AC was reduced by only 29% within 60 minutes for animals treated with *cul-1(RNAi)* (*n* = 31, *P* < 0.0001; **Figure 4D**).

We next wanted to determine if any of the Skp1-related proteins in *C. elegans* function as adaptors that link CUL-1 to the F-box protein TIR1 to mediate degradation of AID-tagged target proteins. Based on the availability of RNAi clones, we fed synchronized L1 stage animals with RNAi targeting four of the eight Skp1-related adaptors known to interact with CUL-1; *skr-1*, *skr-2*, *skr-7*, and *skr-10* (Nayak *et al*. 2002; Yamanaka *et al*. 2002). Owing to the 83% sequence homology between *skr-1* and *skr-2* likely stemming from a gene duplication event and predicted cross-RNAi effects, their gene names are unified in this report as *skr-1/2* similar to (Nayak *et al*. 2002). Of all the Skp1 homologs, *C. elegans skr-1* and human Skp1 share the greatest sequence homology (Yamanaka *et al*. 2002). We also fed animals RNAi targeting *rbx-1*, which encodes the RING finger protein in the SCF E3 ubiquitin ligase (Yamanaka *et al*. 2002). To assess AID::GFP abundance in the AC, we again used animals expressing *eft-*3>AID::GFP and *eft-3>*TIR1::mRuby and examined animals at the P6.p 2-cell stage (**Figure 4B**). RNAi knockdown of *skr-1/2* compared to *control(RNAi)* led to differences in AID::GFP abundance that were not statistically significant (*n* = 23 and 20, respectively, *P* = 0.3522). However, similar to *cul-1(RNAi)*, RNAi silencing of *rbx-1* (*n* = 22) resulted in a statistically significant increase in the abundance of AID::GFP in the AC compared to *control(RNAi)* treatment (*P* = 0.0283) (**Figure 4E**). Interestingly, both *skr-7(RNAi)* (*n* = 23) and *skr-10(RNAi)* (*n* = 21) resulted in a statistically significant decrease in AID::GFP abundance compared to control (*P* < 0.0001).

We also wanted to determine whether depletion of *skr-1/2*, *skr-7*, *skr-10,* and *rbx-1* could inactivate AID-mediated protein degradation We fed synchronized L1 stage animals with RNAi targeting these SCF complex components. We treated animals at the P6.p 2-cell stage with 1 mM NAA for 60 minutes and quantified AID::GFP degradation in the AC (**Figure 4F**). For *control(RNAi)*-treated animals, the abundance of AID::GFP in the AC was once again nearly undetectable within 60 minutes (−96%, *n* = 21) (**Figure 4F**). Similarly, the AID::GFP abundance in animals treated with *skr-7(RNAi)* (*n* = 21) and *skr-10(RNAi)* (*n* = 21) was undetectable within 60 minutes of NAA exposure (−94% and −93%, respectively). For animals treated with *skr-1/2(RNAi)* (*n* = 21, *P* < 0.0001), the abundance of AID::GFP in the AC was reduced by 57% within 60 minutes (**Figure 4F**). For animals treated with *rbx-1(RNAi)* (*n* = 23, *P* < 0.0001), the abundance of AID::GFP in the AC was reduced by 65% within 60 minutes (**Figure 4F**). These results suggest that: 1) suppression of *cul-1*, *skr-1/2*, or *rbx-1* is sufficient to block TIR1-mediated degradation, while suppression of *skr-7* or *skr-10* is not; 2) TIR1 functions as a substrate recognition component of the *C. elegans* CUL-1-based SCF complex, which was also previously shown in yeast (Nishimura *et al*. 2009); and 3) it is possible to deplete multiple targets simultaneously using both AID and RNAi technology.

Inhibiting the expression of *cul-1, skr-1/2, or rbx-1* is a valid approach for reversing AID-mediated degradation in *C. elegans*. We suggest using *cul-1, skr-1/2 or rbx-1(RNAi)* for this purpose with caution, as they have known cell cycle-dependent functions and therefore silencing them may conflate the recovery of AID-tagged proteins with a cell cycle phenotype (Kipreos *et al*. 1996). As an alternative approach to achieving recovery of AID-tagged proteins, we propose the use of RNAi targeting TIR1 or simply using auxinole, a commercially available inhibitor of TIR1 (Hayashi *et al*. 2012; Yesbolatova *et al*. 2019). One caveat to this approach is that auxinole is expensive and thus it may be difficult to obtain stoichiometrically equivalent amounts of auxin and auxinole to truly achieve recovery of one’s protein of interest. However, for *C. elegans* researchers requiring tighter temporal control, these may be avenues worth exploring. Presently, recovery from degradation with 1 mM auxin takes up to 24 hours to fully recover expression of the target protein (Zhang *et al*. 2015). Such protein recovery kinetics are insufficient for studying events in the nematode that occur within minutes to hours such as uterine-vulval attachment, vulval morphogenesis, or many other developmental events occurring post-embryonically.

### NAA as a tool for exploring phenotypes during development and beyond

As our previous results demonstrate that we could effectively deplete a non-functional AID::GFP reporter expressed in the uterine AC and VPCs, we next tested whether NAA-mediated depletion of target proteins could be utilized to study post-embryonic developmental events occurring over a tight temporal window. We focused on a well-studied system of organogenesis, *C. elegans* uterine-vulval cell specification and morphogenesis (**Figure 1B**) (Schindler and Sherwood 2013). As a proof-of-principle, we chose to deplete the nuclear hormone receptor, *nhr-25,* a homolog of arthropod Ftz-F1 and vertebrate SF-1/NR5A1 and LRH-1/NR5A2, which has been shown to function pleiotropically in a wide array of developmental events, from larval molting (Asahina *et al*. 2000; Gissendanner and Sluder 2000; Frand *et al*. 2005), heterochrony (Hada *et al*. 2010), and uterine-vulval morphogenesis (Chen *et al*. 2004; Hwang and Sternberg 2004; Asahina *et al*. 2006; Hwang *et al*. 2007; Ward *et al*. 2013). It was this pleiotropy that made targeting NHR-25 an attractive target, as RNAi and mutant analyses have shown previously that it is initially required in the AC during the AC/VU decision for proper specification of AC fate (Hwang and Sternberg 2004; Asahina *et al*. 2006) and approximately 7 hours later it is required in the underlying VPCs for cell division (Chen *et al*. 2004; Hwang *et al*. 2007; Ward *et al*. 2013).

First, we examined the *nhr-25::GFP::AID* expression pattern, and observed GFP localization to the nuclei of the AC/VU cells during the mid-L2 stage, enrichment in the AC following specification, and nuclear localization in the 1° and 2° VPCs during all stages of vulval division and morphogenesis (**Figure 5A**). We quantified GFP fluorescence over developmental time. Consistent with previous reports based on transgene analyses (Gissendanner and Sluder 2000; Ward *et al*. 2013), endogenous *nhr-25::GFP::AID* AC expression peaks after AC specification in the early L3 at the P6.p 1-cell stage and is undetectable above background by the P6.p 4-cell stage at the time of AC invasion. Conversely, *nhr-25::GFP::AID* increases in intensity in the VPCs at the P6.p 4-cell stage, peaking during the morphogenetic events following AC invasion (**Figure 5B**). Given this temporally-driven expression pattern and based on previous experimental results from RNAi and mutant analyses (Chen *et al*. 2004; Hwang and Sternberg 2004; Asahina *et al*. 2006; Hwang *et al*. 2007; Ward *et al*. 2013), we hypothesized that depleting AID-tagged NHR-25 prior to AC specification should interfere with the AC/VU decision. To test this hypothesis, we used synchronized L1 stage animals expressing *eft-3>*TIR1::mRuby and endogenously tagged NHR-25::GFP::AID. We exposed these larvae to 4 mM NAA or a buffer control and examined animals in the early L3 stage, after the normal time of AC specification. Strikingly, all 36 animals examined showed a failure to specify the AC fate, with the presence of either one (10/36) or two (26/36) small AC/VU-like cells in the central gonad as compared to control animals (**Figure 5C**). Next, we repeated the experiment but waited until after AC specification, in the early L3 stage, to expose animals to buffer control or 4 mM NAA. Here, in all animals, we detected the presence of an AC situated over P6.p, but in 34 of the 36 animals, P6.p failed to divide as compared to controls at the mid-L3 stage (**Figure 5E**). Quantification of *nhr-25::GFP::AID* in AC/VU cells (**Figure 5D**) and VPCs (**Figure 5F**) demonstrated the 4 mM NAA treatment robustly depleted endogenous protein by 95% in the AC/VU and 81% in the VPCs, respectively. Finally, we waited until treated animals (early L3 stage) became adults (approximately 24 hours later) and examined them for plate level phenotypes. We saw a 100% Egg-laying defect (Egl) in 4 mM NAA treated animals as compared to control treated plates (**Figure 5G**). Together, these results indicate that the synthetic auxin, NAA, can robustly deplete target endogenous proteins in a facile, high-throughput fashion during uterine-vulval development. This method should prove valuable in dissecting pleiotropic gene function in the future.

**Figure 5.**
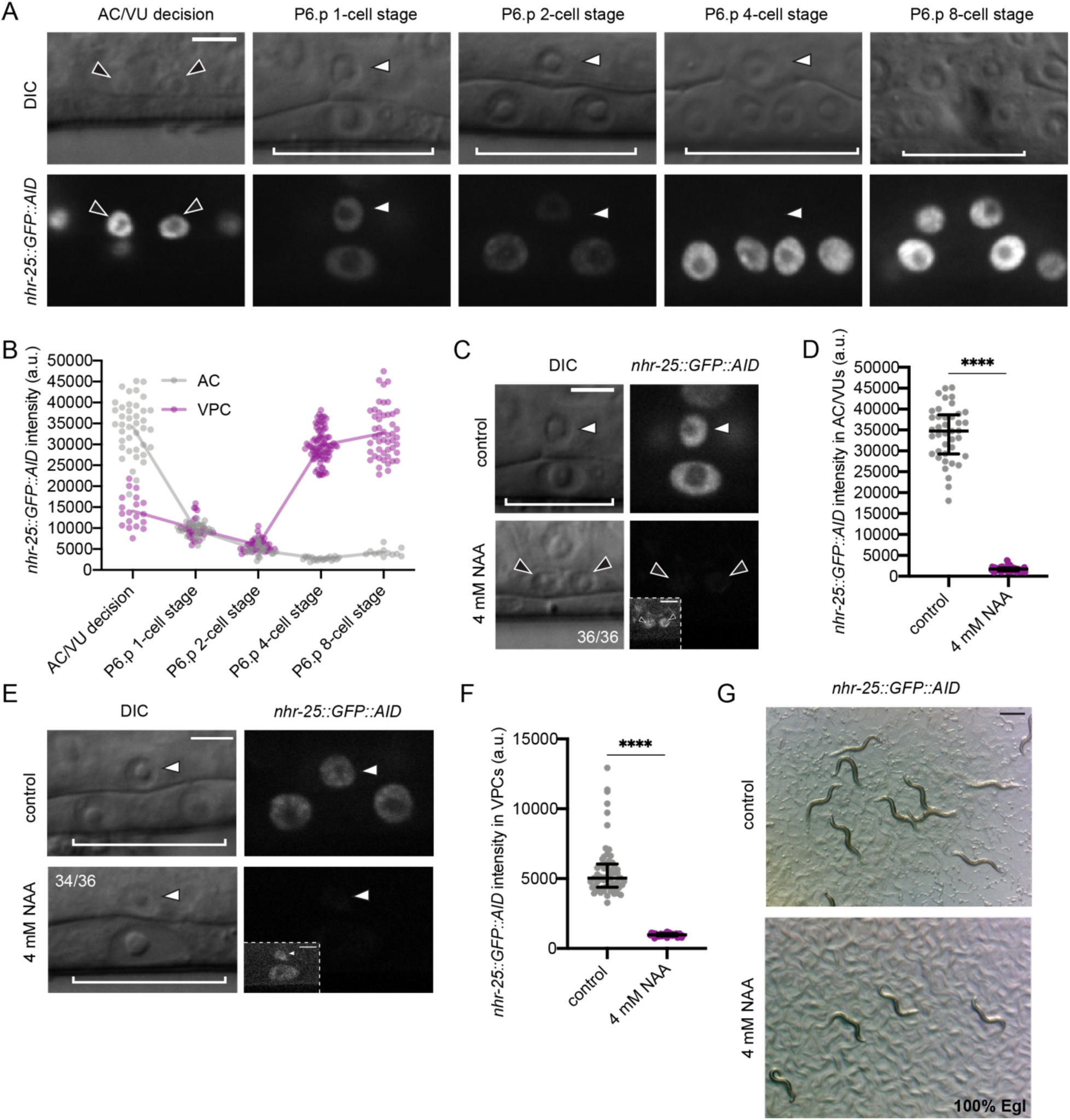
NAA-mediated degradation of NHR-25 causes AC specification and VPC division defects. (A) *nhr-25::GFP::AID* localizes to the nuclei of the AC/VU (black arrowheads), the AC (white arrowheads) and VPCs (brackets). At P6.p 8-cell stage (far right) the AC is not in the same focal plane as the 1° VPCs. (B) Quantification of *nhr-25::GFP::AID* over developmental time, from the AC/VU decision to the P6.p 8-cell stage. The curve is connected by the mean at each developmental stage (*n* = 20, 31, 20, 21, and 12 animals quantified, respectively). (C) DIC and corresponding GFP images of ACs (arrowheads) and underlying VPCs (brackets) from early L3 stage animals. Animals expressing *nhr-25::AID::GFP* and *eft-3>*TIR1::mRuby were treated with control and 4 mM NAA. (D) Quantification of *nhr-25::GFP::AID* in AC/VUs following NAA treatment. Data presented as the median+IQR (*n* ≥ 20 animals examined for each, and *P* < 0.0001 by a Mann Whitney U test). (E) DIC and corresponding GFP images of ACs (arrowheads) and underlying VPCs (brackets) from mid-L3 stage animals. Animals expressing *nhr-25::AID::GFP* and *eft-3>*TIR1::mRuby were treated with control and 4 mM NAA. (F) Quantification of *nhr-25::GFP::AID* in VPCs following NAA treatment. Data presented as the median+IQR (*n* ≥ 30 animals examined for each, and *P* < 0.0001 by a Mann Whitney U test). (G) Representative images of adult plate level phenotypes following control and 4 mM NAA treatments added at the L3 stage (*n* ≥ 30 animals examined). Scale bar in (G), 500 µm.

### Important caveats of the C. elegans AID system

Our results reported here with a synthetic auxin, NAA, as well as the initial report developing the *C. elegans* AID system (Zhang *et al*. 2015) clearly demonstrate the effectiveness of auxin-induced targeted protein depletion. Several other recent reports have also effectively used the AID system in *C. elegans* to control protein function, including controlling spermatogenesis by manipulating *spe-44* levels (Kasimatis *et al*. 2018), depleting a mediator component to modulate longevity (Lee *et al*. 2019), examining chromosome segregation during oogenesis (Ferrandiz *et al*. 2018), meiotic crossover (Zhang *et al*. 2018), and revealing novel roles of neuronal gene function through conditional depletion (Serrano-Saiz *et al*. 2018). Despite the increasing frequency of AID system usage in the *C. elegans* community, there are only a handful of TIR1 driver lines published, and the importance of copy number and promoter strength has not been systematically assessed.

While we are optimistic that the use of the synthetic analog of auxin presented here will allow for even more widespread utility of the AID system in the *C. elegans* research community, there are still are some areas open to improvement for the technology. A recent report in mammalian cell culture identified that AID-tagged proteins are depleted in an auxin-independent fashion in the presence of TIR1, relative to wild-type levels (Li *et al*. 2019; Sathyan *et al*. 2019). We examined if this was also occurring in *C. elegans* strains in our laboratory expressing AID-tagged proteins and TIR1. We were able to detect statistically significant auxin-independent depletion of both a ubiquitously expressed AID::GFP transgene under the *eft-3* promoter in ACs (−22%, *n* = 24, *P* < 0.0001) and VPCs (−24%, *n* = 24, *P* < 0.0001; **Figure S4A-B**) and an endogenously tagged *nhr-25::GFP::AID* allele in ACs (−22%, *n* = 26, *P* = 0.0022) and VPCs (−35%, *n* = 26, *P* < 0.0001; **Figure S4C-D**). As partial loss of an endogenous protein could generate a hypomorphic condition, both from the placement of the AID tag and apparent triggering of the degradation machinery, we urge caution in carefully evaluating AID-tagged alleles paired with TIR1, independent of auxin delivery. Further optimization of the AID system in *C. elegans* will hopefully ameliorate this concern, as researchers recently used the heterologous co-expression of an auxin response factor (ARF) with TIR1 to rescue auxin-independent degradation in mammalian cell culture (Sathyan *et al*. 2019).

### Conclusion

The ease of editing the *C. elegans* genome using CRISPR/Cas9-based approaches (Calarco and Friedland 2015) and heterologous gene manipulation tools is ushering in a new era of cellular and developmental biology. Several new tools available to *C. elegans* researchers require the insertion of small amino acid tags into target loci, including ZF1 tagging (Armenti *et al*. 2014), sortase A (Wu *et al*. 2017), and the AID system (Zhang *et al*. 2015). Alternatively, any GFP fusion can be targeted via a GFP nanobody tethered to ZIF1 (Wang *et al*. 2017). These genomic edits are then paired with single transgene expression to allow for targeted spatial and temporal loss-of-function approaches through manipulation of endogenous loci. Prior to their advent, spatial and temporal control of protein function was largely missing from the *C. elegans* genomic toolkit. With an ever-increasing set of these tools being optimized for *C. elegans*, it is clear that different tools will have strengths and weaknesses depending on multiple variables, including subcellular localization of target protein, availability of tissue- and cell-type specific drivers, and inducibility of depletion. Here, we optimize a powerful heterologous system, the auxin-inducible degradation system. We demonstrate that a synthetic auxin analog, NAA, and its water-soluble, potassium salt, K-NAA, can function equivalently to natural auxin. The water solubility permits easier preparation of media and allows researchers to perform experiments in liquid culture and microfluidics. Importantly, the use of ethanol free K-NAA may be beneficial to *C. elegans* researchers studying behavior and aging, where introduction of ethanol may lead to confounding results. We also demonstrate the strength of the AID system for studying developmental cell biology by examining multiple spatial and temporal roles of the Ftz-F1 homolog *nhr-25* during uterine and vulval morphogenesis. It is our hope that the use of the synthetic auxin NAA will complement the AID system in *C. elegans* when examining targeted protein depletion phenotypes in tissues and developmental stages of interest. As the library of tissue-specific TIR1 drivers continues to grow, we envision researchers being able to rapidly degrade proteins of interest in specific tissues and visualize the outcome at single-cell resolution.

## ACKNOWLEDGEMENTS

We thank R. Adikes for the suggestion to look into a water-soluble form of auxin and B. Lacroix of the Vitaly Citovsky lab for providing the initial aliquot of NAA. We thank R. Adikes, J. Smith and N. Palmisano for insightful discussions while developing the liquid based NAA protocol and R. Adikes for helpful advice regarding image acquisition parameters and quantification. We thank N. Stec of the Christopher Hammell lab for help with the microfluidic device. We thank M. Li and N. Kim of the Jessica Seeliger lab for help with the *E. coli* OP50 growth curve. We thank R. Adikes, N. Palmisano, A. Kohrman and A. Nishimura for advice and comments on the manuscript. We appreciate the help of W. Zhang for preparing media and plates.

This work was funded by the National Institute of Health (NIH) National Institute of General Medical Sciences (NIGMS) [1R01GM121597-01 to D.Q.M.]. D.Q.M. is also a Damon Runyon-Rachleff Innovator supported (in part) by the Damon Runyon Cancer Research Foundation [DRR-47-17]. M.A.Q.M. is supported by the NIH NIGMS [2T32GM007518-41] and [3R01GM121597-02S2]. T.N.M. is supported by the NIH NICHD [1F31HD100091-01]. C.M.H and B.A.K. are supported by the NIH NIGMS [R01GM117406]. J.D.W. is supported by the NIH NIGMS [R00GM-107345]. Some strains were provided by the *Caenorhabditis* Genetics Center, which is funded by the NIH Office of Research Infrastructure Programs [P40 OD010440].

## AUTHOR CONTRIBUTIONS

M.A.Q.M. and D.Q.M conceived and designed the experiments. G.A. and J.D.W. designed the constructs. J.M.R. performed the microinjections. G.A. performed the crosses and characterized strains. M.A.Q.M, B.A.K., T.N.M., L.J. and J.A. performed the experiments. M.A.Q.M. and D.Q.M. analyzed and quantified the data. M.A.Q.M. and D.Q.M. wrote the manuscript with contributions from the other authors. The authors declare no competing interests.

