## Supplemental Materials for "Rapid degradation of *C. elegans* proteins at single-cell resolution with a synthetic auxin"

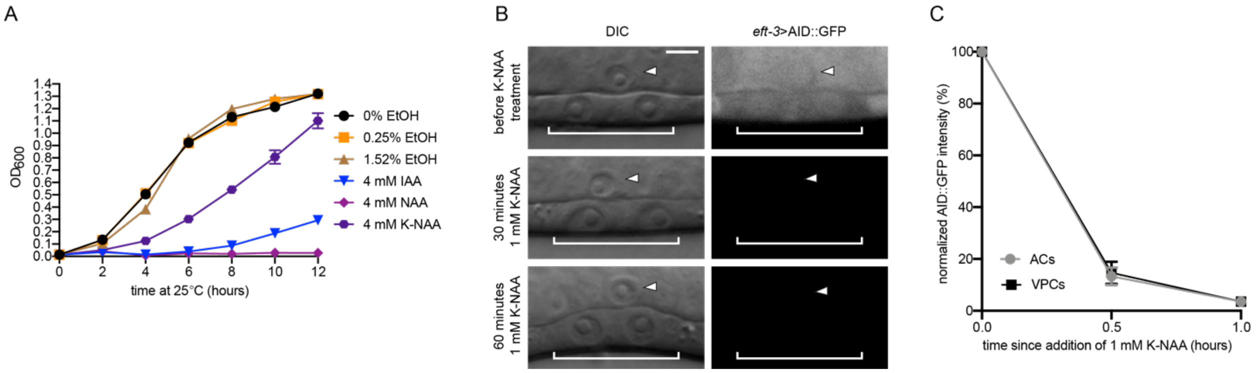


**Figure S1. Degradation kinetics of K-NAA and the effect of auxin on bacterial growth.** (A) OD600 growth curves for *E. coli* OP50 exposed to different percentages of ethanol (0%, 0.25%, and 1.52%) and different forms of auxin (IAA, NAA, and K-NAA). (B) DIC and corresponding GFP images of ACs (arrowheads) and underlying VPCs (brackets) from mid L3 stage animals at the P6.p 2-cell stage. Animals expressing *eft-3*>AID::GFP and *eft-3*>TIR1::mRuby were imaged before (top), 30 minutes (middle) and 60 minutes (bottom) after 1 mM K-NAA exposure. (C) Rates of degradation determined by quantifying AID::GFP in ACs (grey circles) and (VPCs (black squares) following K-NAA treatment. Data presented as the mean±SD ($n \geq 26$ animals examined for each time point).

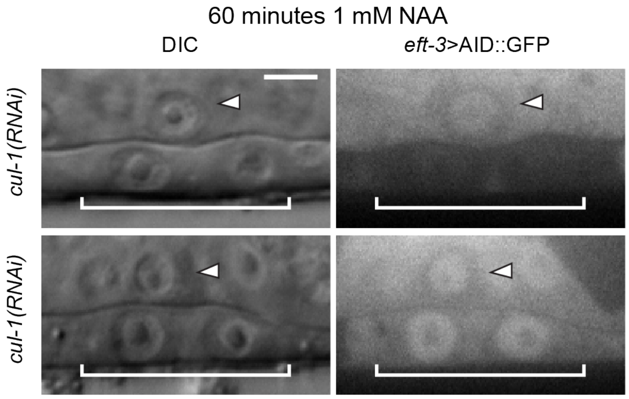


**Figure S2. VPCs are variably sensitive to RNAi.** DIC and corresponding GFP images of ACs (arrowheads) and underlying VPCs (brackets) from mid L3 stage animals at the P6.p 2-cell stage, showing insensitivity to RNAi in the VPCs (top, right) as compared to sensitivity to RNAi (bottom, right). Synchronized L1 stage animals expressing *eft*-3>AID::GFP and *eft*-3>TIR1::mRuby were fed *cul-1(RNAi)* and treated with NAA at the P6.p 2-cell stage.

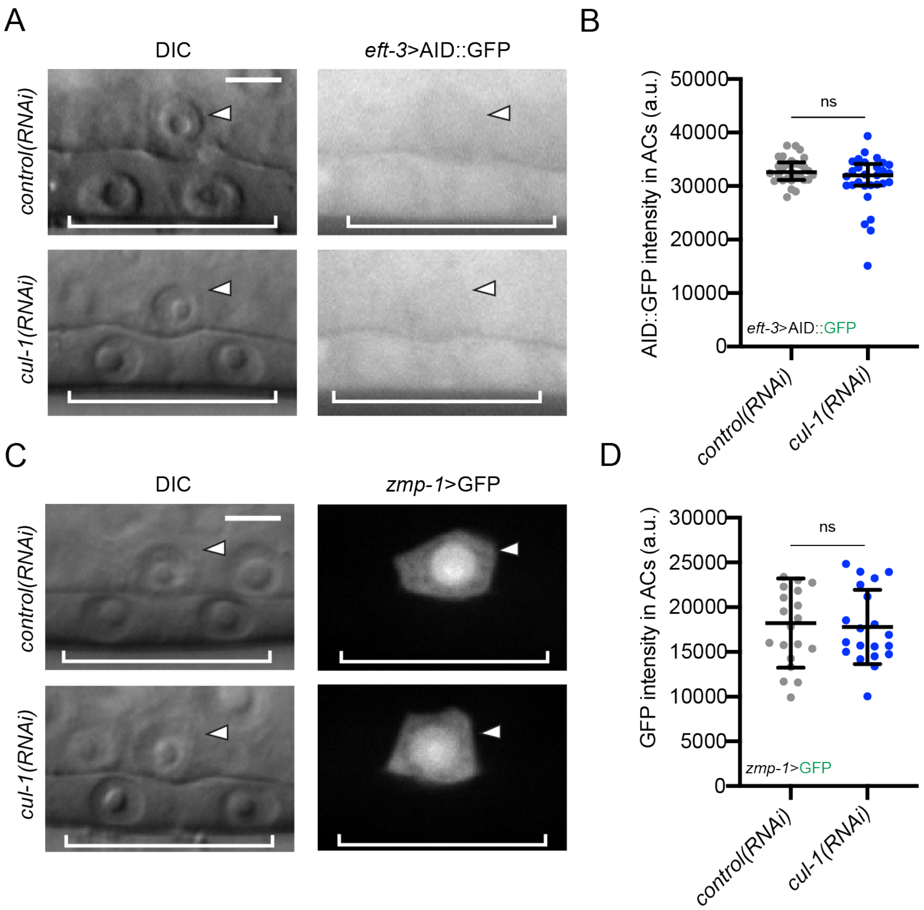


**Figure S3. *cul-1(RNAi)* fails to increase the expression of AID-tagged transgenes in the absence of TIR1.** (A) DIC and corresponding GFP images of ACs (arrowheads) and underlying VPCs (brackets) from mid L3 stage animals at the P6.p 2-cell stage. Animals expressing *eft*-3>AID::GFP without TIR1 in the background were treated with *cul-1(RNAi)*. (B) Quantification of *eft*-3>AID::GFP in ACs. Data presented as the mean±IQR (*n* = 29 animals examined for each, and *P* = 0.1168 by a Mann Whitney U test). (C) DIC and corresponding GFP images of ACs (arrowheads) and underlying VPCs (brackets) from mid L3 stage animals at the P6.p 2-cell stage. Animals expressing *zmp-1*>GFP without TIR1 in the background were treated with *cul-1(RNAi)*. (D) Quantification of *zmp-1*>GFP in ACs. Data presented as the mean±SD (*n* = 20 animals examined for each, and *P* = 0.7682 by a Student's t-test).

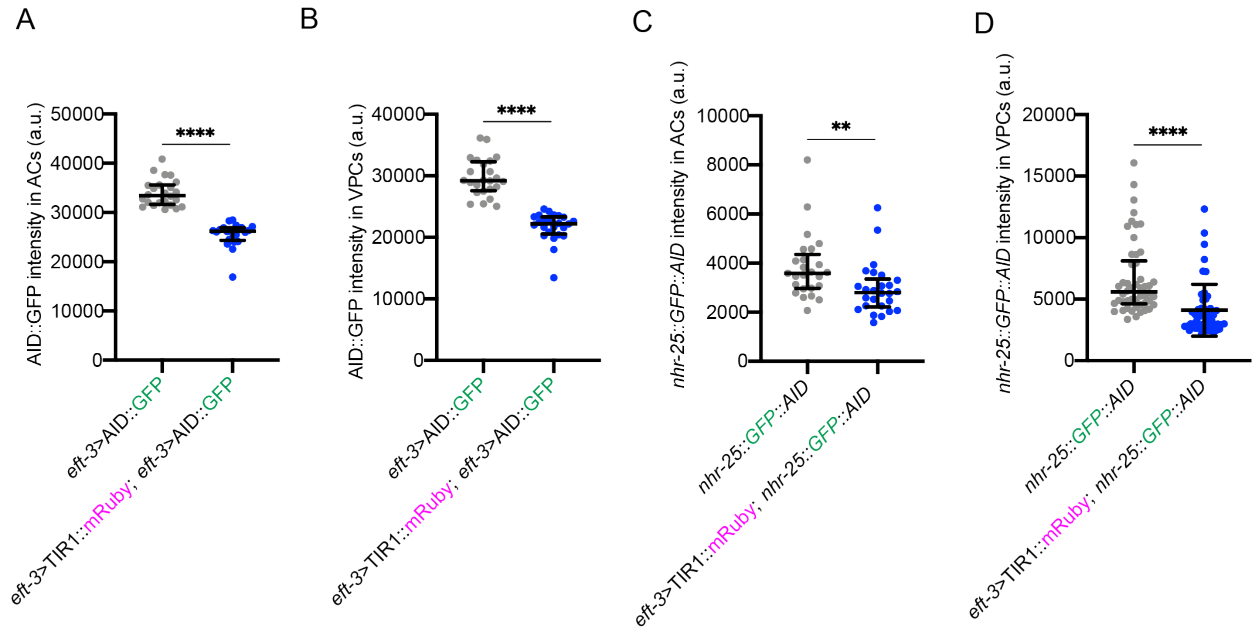


**Figure S4. Auxin-independent degradation of *eft-3*>AID::GFP and *nhr-25::GFP::AID*.** (A-B) Quantification of AID::GFP in (A) ACs and (B) VPCs in a genetic background without (left, grey) and with *eft-3*>TIR1::mRuby (right, blue). Data presented as the median±IQR ($n \geq 24$ animals examined for each, and $P < 0.0001$ by a Mann Whitney U test). (C-D) Quantification of *nhr-25::GFP::AID* in (A) ACs and (B) VPCs in a genetic background without TIR1 (left, grey) and with TIR1 (right, blue). Data presented as the median±IQR ($n \geq 25$ animals examined for each, and $P = 0.0022$ and $P < 0.0001$, respectively by a Mann Whitney U test).

**Table S1. Brood size and embryonic lethality in animals treated with K-NAA, IAA and their respective controls.**

| STRAIN | TREATMENT (25°C) | BROOD SIZE (±SD) | % EMBRYONIC LETHALITY (±SD) |
|---|---|---|---|
| *eft-3*>TIR1::mRuby; *nhr-23::AID* | 0% EtOH | 220.0 (32.9) | 2.0 (1.3) |
| *eft-3*>TIR1::mRuby; *nhr-23::AID* | 4 mM K-NAA | 219.5 (40.1) | 1.88 (0.68) |
| *eft-3*>TIR1::mRuby; *nhr-23::AID* | 0.25% EtOH | 208.3 (41.1) | 2.6 (1.3) |
| *eft-3*>TIR1::mRuby; *nhr-23::AID* | 4 mM IAA | 222.3 (29.8) | 3.1 (1.1) |

**File S1. Protocol for liquid based NAA-mediated degradation in *C. elegans*.**

| STEP | SOLUTION | COMMENTS |
|---|---|---|
| (1) Wash off synchronized animals with 500 µL of NAA in M9 from an NGM plate seeded with *E. coli* OP50. | Dilute NAA (N1641) in 1X M9 buffer to the desired concentration. | NAA arrives at 5.4 mM. Store at 4°C. |
| (2) Transfer animals to a concave well on a spot plate. | N/A | Transferring animals to a microcentrifuge tube is not recommended, as animals need to thrash freely. |
| (3) House the spot plate inside a homemade humidity chamber. | N/A | An empty pipet box suffices as a homemade humidity chamber. Place wet paper towels along the edges of the box and place dry paper towels along the edges of the lid. |
| (4) Seal the pipet box with parafilm. | N/A | N/A |
| (5) Incubate for a desired period of time. | N/A | N/A |
| (6) Take up 1.2 µL from the bottom of the well. | N/A | N/A |
| (7) Dispense onto a 5% agar pad containing 10 mM sodium azide and secure animals with a coverslip. | N/A | N/A |
| (8) Image animals under a microscope. | N/A | N/A |

**File S2. Protocol for the preparation of synthetic auxin-containing media for the *C. elegans* larvae-specific microfluidic device (Keil *et al.* 2017).**

| STEP | SOLUTION | COMMENTS |
|---|---|---|
| (1) Pick single colony of *E. coli* NA22 to 1 L of LB. | N/A | N/A |
| (2) Incubate and shake at 37°C for approximately 20 hours. | N/A | N/A |
| (3) Spin for 20 minutes at 2000 x g. | N/A | N/A |
| (4) Remove the supernatant. | N/A | N/A |
| (5) Dilute the pellet in 60 mL of M9 buffer and transfer to two 50 mL conical centrifuge tubes. | N/A | Filter sterilize M9 buffer before use. |
| (6) Repeat steps 4 and 5 two more times. | N/A | N/A |
| (7) Resuspend the final pellet in each conical tube in 25 mL of 4 mM K-NAA in M9 buffer. | Dilute K-NAA (#N610) in 1X M9 buffer to a concentration of 4 mM. | Store at 4°C. |
| (8) Dilute the bacterial culture to an $OD_{600}$ equal to 7. | N/A | To achieve an $OD_{600}$ equal to 7, dilute 1:10 with 4 mM K-NAA in M9 buffer. |
| (9) Run media through the microfluidic device. | N/A | Store the media at 4°C for up to 2 weeks. |

**Reagent Table**

| REAGENT | SOURCE | IDENTIFIER |
|---|---|---|
| **Bacterial and Virus Strains** | | |
| *E. coli*: Strain OP50 | Caenorhabditis Genetics Center | WormBase ID: OP50 |
| *E. coli*: Strain NA22 | Caenorhabditis Genetics Center | WormBase ID: NA22 |
| **Chemicals, Peptides, and Recombinant Proteins** | | |
| Indole-3-acetic acid, 98+% | Alfa Aesar | A10556 |
| 1-Naphthaleneacetic acid | Sigma-Aldrich | 317918 |
| 1-Naphthylacetic acid | Sigma-Aldrich | N1641 |
| Naphthaleneacetic acid (K-NAA) | PhytoTechnology Laboratories | N610 |
| NEBuilder HiFi DNA Assembly Master Mix | New England Biolabs | E2621 |
| PureLink HQ Mini Plasmid DNA Purification Kit | Invitrogen | K210001 |
| **Experimental Models: Organisms/Strains** | | |
| *C. elegans*: Strain CA1202: *ieSi57 [eft-3>TIR1::mRuby::unc-54 3'UTR + Cbr-unc-119(+)] II; ieSi58 [eft-3>degron::GFP::unc-54 3'UTR + Cbr-unc-119(+)] IV* | Caenorhabditis Genetics Center | WormBase ID: CA1202 |
| *C. elegans*: Strain CA1204: *ieSi58 [eft-3>degron::GFP::unc-54 3'UTR + Cbr-unc-119(+)] IV* | Caenorhabditis Genetics Center | WormBase ID: CA1204 |
| *C. elegans*: Strain JDW58: *nhr-25(wrd11[nhr-25::GFP^SEC^degron:3xFLAG]) X* | This paper | N/A |
| *C. elegans*: Strain JDW59: *nhr-25(wrd12[nhr-25::GFP^degron:3xFLAG]) X* | This paper | N/A |
| *C. elegans*: Strain JDW70: *nhr-25(wrd11[nhr-25::GFP^SEC^degron:3xFLAG]) X; unc-119(ed3); ieSi57* | This paper | N/A |
| *C. elegans*: Strain JDW71: *nhr-25(wrd18[nhr-25::GFP^degron:3xFLAG]) X; unc-119(ed3); ieSi57* | This paper | N/A |
| *C. elegans*: Strain KRY88: *nhr-23(kry61[nhr-23::AID-TEV-3xFLAG]) I*; *ieSi57 [eft-3>TIR1::mRuby::unc-54 3'UTR + Cbr-unc-119(+)] II* | Zhang *et al.*, 2015 | N/A |
| *C. elegans*: Strain PS3239: *syIs49 [zmp-1>GFP + (pMH86) dpy-20(+)] IV* | Caenorhabditis Genetics Center | WormBase ID: PS3239 |
| **Oligonucleotides** | | |
| 30x linker for CT slot-F (#3775) CTGTGGCTCTTCGGCGGGTGGCGGTGGATCGG | This paper | N/A |
| 30x linker for CT slot-R (#3776) AAACGCTCTTCGCATGGATCCGCCTCCTCCTGAAC | This paper | N/A |
| *nhr-25* 5'homology arm-F (#3478) GGCTGCTCTTCGTGGAGTGCTCATCTCTCCCTCCA | This paper | N/A |
| *nhr-25* 5'homology arm-R (#3479) GGGTGCTCTTCGCGCTGATGCCATATAAGGTACTG CGGTATATGTGGCTTGTGGGGTTGGC | This paper | N/A |
| attB1/B2+SapI extension for Gateway cloning out 5' homology arms-F (#3631) GGGGACAAGTTTGTACAAAAAAGCAGGCTGGCTGG CTGCTCTTCGTGG | This paper | N/A |

| | | |
|---|---|---|
| attB1/B2+SapI extension for Gateway cloning out 5' homology arms-R (#3632)<br>GGGGACCACTTTGTACAAGAAAGCTGGGTCGCTCT TCGCGC | This paper | N/A |
| Amplify *nhr-25* 3’ homology arm by TOPO blunt cloning (#3480)<br>GGCTGCTCTTCGACGTAATGTCTCTGTCATAGGAGC TGAAAAC | This paper | N/A |
| Amplify *nhr-25* 3’ homology arm by TOPO blunt cloning (#3481)<br>GGGTGCTCTTCGTACAGAAGAGTTCGGCAGAGAAC G | This paper | N/A |
| *nhr-25* genotyping (anneals outside homology arms; #3954)<br>TGGACTCTTTCCCAGCAGTC | This paper | N/A |
| *nhr-25* genotyping (anneals outside homology arms; #3955)<br>GGCGAGGAAAATAAAGGGCAAA | This paper | N/A |
| Degron-specific primer for sequencing out of 3’ end of the knock-in (#3380)<br>AAGAACGTGATGGTTTCCTGC | This paper | N/A |
| GFP-specific primer for sequencing out of 5’ end of knock-in (#3378)<br>TGGACTCCGTAGCAGAAGGTGG | This paper | N/A |
| *nhr-25* genotyping for crosses (following knock-in confirmation; #1584)<br>AGAGAAGAGAAGCATCGGAAG | This paper | N/A |
| *nhr-25* genotyping for crosses (following knock-in confirmation; #1585)<br>TGTGAGGGTTTGGGCACTAGG | This paper | N/A |
| CA1200 genotyping (oCFJ1493)<br>TGCTCGGAAGGACTTGATTT | Frøkjær-Jensen *et al.*, 2014 | N/A |
| CA1200 genotyping (oCFJ1494)<br>TTGCCACGTCTTCTTGAGTG | Frøkjær-Jensen *et al.*, 2014 | N/A |
| CA1200 genotyping (oCFJ1529)<br>TATCGTAAATCGGCGCGAGC | Frøkjær-Jensen *et al.*, 2014 | N/A |
| *nhr-25* sgRNA-F (#3482)<br>TTGATACACTGCTGTGCCGTACA | This paper | N/A |
| *nhr-25* sgRNA-F (#3482)<br>AACTGTACGGCACAGCAGTGTAT | This paper | N/A |
| **Recombinant DNA** | | |
| 30xlinker gBlock<br>CTTTCCTGCGTTATCCCCTGATTCTGTGGCTCTTCg GCGGGTGGCGGTGGATCGGGAGGAGGAGGTTCGG GTGGCGGAGGCAGTGGAGGTGGCGGAAGTGGCGG CGGTGGTTCAGGAGGAGGCGGATCCATGcGAAGAG CGTTTGATGCCTGGCAGTTTATGGCGGG | This paper; IDT | |

| | | |
|---|---|---|
| BglI-cul-1-SalI_T444T gBlock<br>CAAGGGGTTATGCTAGTAATACGACTCACTATAGGG<br>AGACCGGCAGAAGACGGTCGAGCAGAATCTTCAAC<br>TCCAGCACGAACCGCTGGCGCGGATTTTGTCGGTC<br>ACGAGATGTACCAGCGTGTTGAGGAATATGTAAAAG<br>CGTACGTAATCGCTGTTTGTGAAAAGGGCGCCGAG<br>TTGTCGGGCGAGGATCTTCTGAAATACTACACAACT<br>GAATGGGAGAACTTCCGAATCTCGTCGAAGGTGAT<br>GGATGGTATTTTTGCCTACCTCAATCGTCATTGGAT<br>CAGACGTGAGCTCGACGAGGGCCATGAGAACATTT<br>ACATGGTTTATACACTTGCGTTGGTCGTCTGGAAGC<br>GTAACTTGTTCAATGATCTTAAGGATAAGGTGATTG<br>ATGCGATGCTCGAGCTGATTCGTTCCGAGAGAACC<br>GGATCCATGATCAACAGCCGATACATTTCTGGAGTT<br>GTCGAGTGTCTTGTGGAGTTGGGTGTTGACGATTC<br>GGAGACCGATGCGAAGAAGGACGCGGAGACCAAG<br>AAGCTGGCCGTTTACAAGGAATTCTTCGAAGTCAAG<br>TTTTTGGAGGCGACCCGCGGATTTTATACCCAAGAG<br>GCGGCAAATTTCCTGAGCAACGGAGGAAATGTGAC<br>AGATTACATGATTAAAGTGGAGACAAGACTTAATCA<br>AGAGGATGATCGCTGTCAACTCTACCTGAATTCTTC<br>CACAAAAACTCCACTTGCAACTTGCTGTGAATCTGT<br>GCTCATCTCTAATCAGCTTGACTTCCTCCAGAGACA<br>CTTCGGTGGTCTTCTTGTCGATAAACGAGACGATGA<br>CCTTTCCAGAATGTTCAAGCTTTGCGATAGAGTGCC<br>GAATGGACTCGACGAGCTCAGAAAATCCCTGGAGA<br>ATCACATCGCAAAGGAGGGTCACCAGGCGTTGGAA<br>AGAGTTGCTATGGAGGCGGCAACCGACGCGAAACT<br>CTACGTGAAGACGCTTCTGGAGGTTCATGAACGTTA<br>TCAAAGCCTTGTCAATCGATCATTCAAAAATGAGCC<br>AGGATTCATGCTCGACCTCGAGGGGGGGCCCG | This paper; IDT | |
| **Software and Algorithms** | | |
| Fiji | Schindelin et al., 2012 | Version: 2.0.0-rc-69/1.52p |
| Prism | GraphPad | Version: 8.1.2 |
| BioRender | BioRender | biorender.com |
| ChemDraw | PerkinElmer | Version: 18.0 |
| MetaMorph | Molecular Devices | Version: 7.8.12.0 |
| Photoshop | Adobe | Version: 20.0.6 |
| Illustrator | Adobe | Version 23.0.26 |
